# Pred-AHCP: Robust feature selection enabled Sequence-Specific Prediction of Anti-Hepatitis C Peptides via Machine Learning

**DOI:** 10.1101/2024.05.05.592323

**Authors:** Akash Saraswat, Utsav Sharma, Aryan Gandotra, Lakshit Wasan, Sainithin Artham, Arijit Maitra, Bipin Singh

## Abstract

Every year, an estimated 1.5 million people worldwide contract Hepatitis C (HepC), a significant contributor to liver disease. Although many studies have explored machine learning’s potential to predict antiviral peptides, very few have addressed predicting peptides against specific viruses such as Hepatitis C. In this study, we demonstrate the use of machine learning (ML) algorithms to predict peptides that are effective against HepC. We developed an explainable ML model that harnesses the amino acid sequence of a peptide to predict its potential as an anti-HepC (AHC) agent. Specifically, features were computed based on sequence and physicochemical properties, with feature selection performed utilizing a combined scheme of mutual information and variance inflation factor. This facilitated the removal of redundant and multicollinear features from the sequence data, enhancing the model’s generalizability in predicting AHCPs. The model using the *random forest* algorithm produced the best performance with an accuracy of about 90%. The feature selection analysis highlights that the distribution of hydrophobicity and polarizability, as well as the frequencies of glycine residues and di-peptide motifs—YXL, LXK, VXXXF, VL, LV, CC, RR, TXXXV, VXXA, CXXXC—emerged as the key predictors for identifying AHCPs targeting different components of the HepC virus. The model developed can be accessed through the Pred-AHCP web server, provided at http://tinyurl.com/web-Pred-AHCP. This resource facilitates the prediction and re-engineering of AHCPs for designing peptide-based therapeutics while also proposing an exploration of similar strategies for designing peptide inhibitors effective against other viruses.

## 1. INTRODUCTION

Hepatitis C virus (HCV) infects the liver, causing inflammation. There are two major categories of HCV infection—*acute* and *chronic*—based on the duration of the infection. Asymptomatic infections characterise acute hepatitis and are not life-threatening. It represents approximately 30% of total infections; generally, virus particles are cleared from the system within six months. However, long exposures develop chronic HCV infections, representing the majority of symptomatic infections. This usually increases the risk of liver cirrhosis and hepatocellular carcinoma [1]. HCV generally spreads through blood transfusion from an infected person. There are no vaccines available for HCV, and existing drugs face challenges of adverse effects, drug resistance, and drug-drug interactions. Peptides have several advantages over conventional small molecules, such as higher selectivity, specificity, solubility, low toxicity, tunable membrane permeability and easy synthesizability [2]. Experimental methods to screen and identify anti-Hepatitis C peptides (AHCPs) are burdensome, consuming both time and resources and underscoring the need for reliable and accurate computational approaches to predicting AHCPs. Short peptides have demonstrated significant potential in targeting various components of different viruses, including HCV, in clinical and preclinical studies [3]. Considerable efforts have been devoted to predicting antimicrobial and antiviral peptides utilising machine learning techniques, focusing on integrating a wide range of features derived from peptide sequences to develop predictive models. Notably, numerous sequence-based profiles of features comprising amino acid composition, dipeptide and tripeptide composition, positional indicators, and physicochemical information are frequently applied in bioactive peptide prediction [4,5].

Previous literature has reported extensively on feature descriptors derived from peptide sequences [6–9]. Nonetheless, the complexity of these features often results in a somewhat limited intuitive understanding of data-driven models [10]. We used a two-fold strategy utilizing mutual information and variance inflation factor to filter specific features that contribute the most in classifying AHCP activity by removing redundant and multicollinear features. In this work, ML models were developed using a manually curated AHCP sequence dataset and a negative class peptide sequence dataset. Figure 1 provides a schematic representation of the model development process. Employing features derived from these peptide datasets, several binary classification models, including a random forest model, were created for AHCP prediction. The random forest model with the best classification accuracy for AHCPs can be accessed as an online web server at http://tinyurl.com/web-Pred-AHCP for screening and re-engineering purposes.

**Figure 1.**
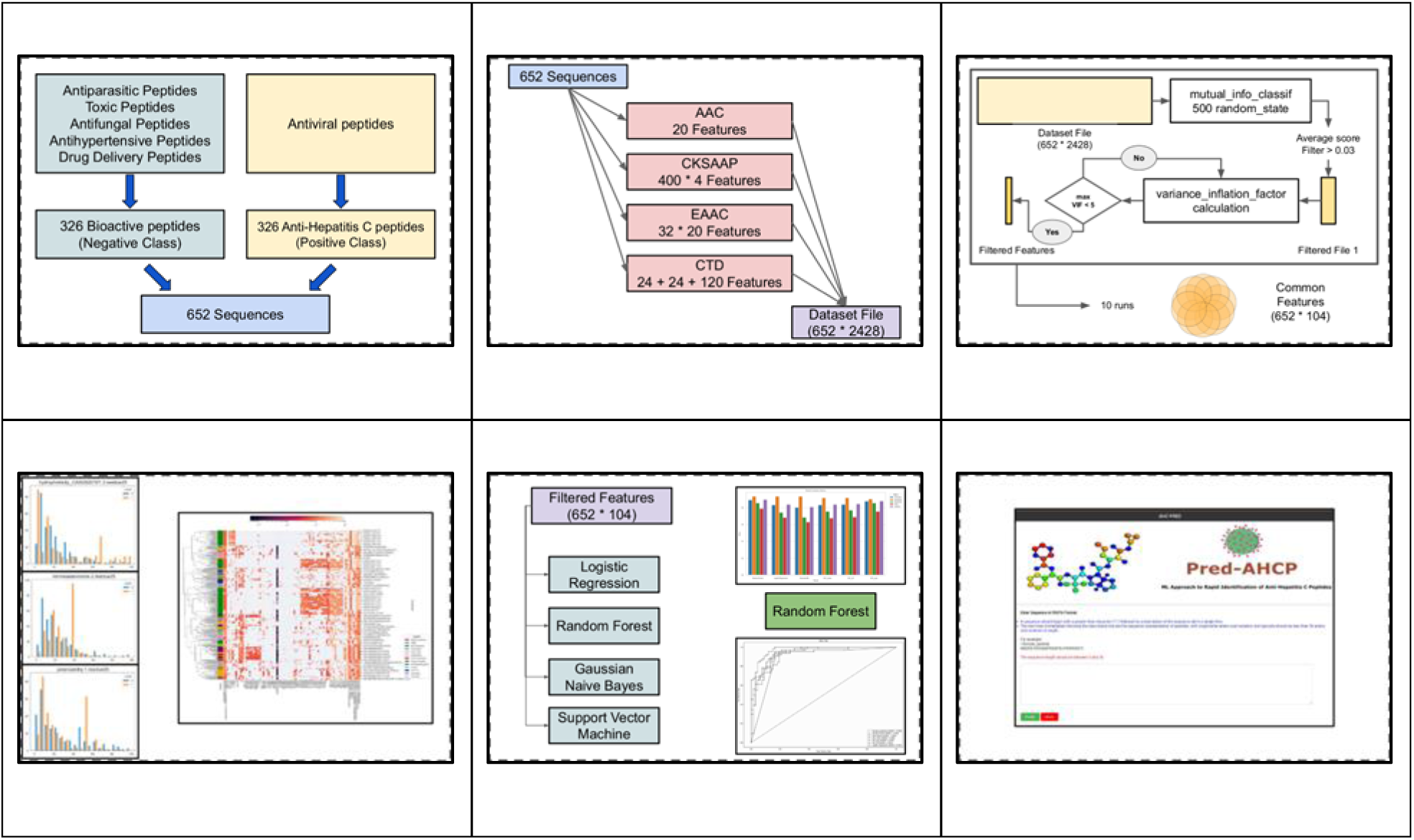
Schematic framework of the Pred-AHCP model. Essential steps are highlighted starting with (a) data collection and validation, (b) feature calculation, (c) selection of key features, (d) feature importance analysis, (e) ML model building, and (f) web server development.

## 2. RESULTS AND DISCUSSION

### 2.1 Selection of Features

A range of diverse global and local sequence-based features was computed on a peptide sequence dataset consisting of benchmarked AHCPs [11] and non-AHCPs (negative dataset), carefully selected based on the length and amino acid composition distribution. Refer to the Materials and Methods section for more details. These feature vectors capture the composition and distribution of amino acids and their physicochemical properties, such as hydrophobicity, charge, polarity, polarizability, secondary structure, solvent accessibility and van der Waals volume of amino acids [12]. Given the high dimensional nature of these features, a two-fold selection strategy was employed to ensure that only crucial and non-redundant features pertinent to AHCPs were incorporated for classification model development. For this, mutual information (MI) was calculated using the *sklearn* (python) library: ‘mutual_info_classif’ [13–15]. To address a feature’s rank variance, the MI score for a feature was obtained by averaging its MI scores estimated from 500 independent trials using different random states to determine the relevant features. To demonstrate the details of filtering criteria, we show a small subset of calculations with 50 realisations of random state. Figure 2 describes the feature indices along the x-axis, arranged according to their mean MI values (y-axis). The dots represent all the MI values of an attribute from the 50 realisations. The blue line highlights the mean or expectation <u>*MI*</u> value of each feature, arranged in descending order, and the red line indicates the threshold considered for attribute filtering. As evident from Figure 2, a sharp drop in MI was observed in the interval 0.03 < <u>*MI*</u> < 0.22. Hence, features with an <u>*MI*</u> value over 0.03 were selected for further analysis and modelling. The trend also highlights that most features have MI values close to or equal to zero, implicating the presence of redundant and irrelevant features that lack the explanatory capability to predict bioactivity class.

**Figure 2.**
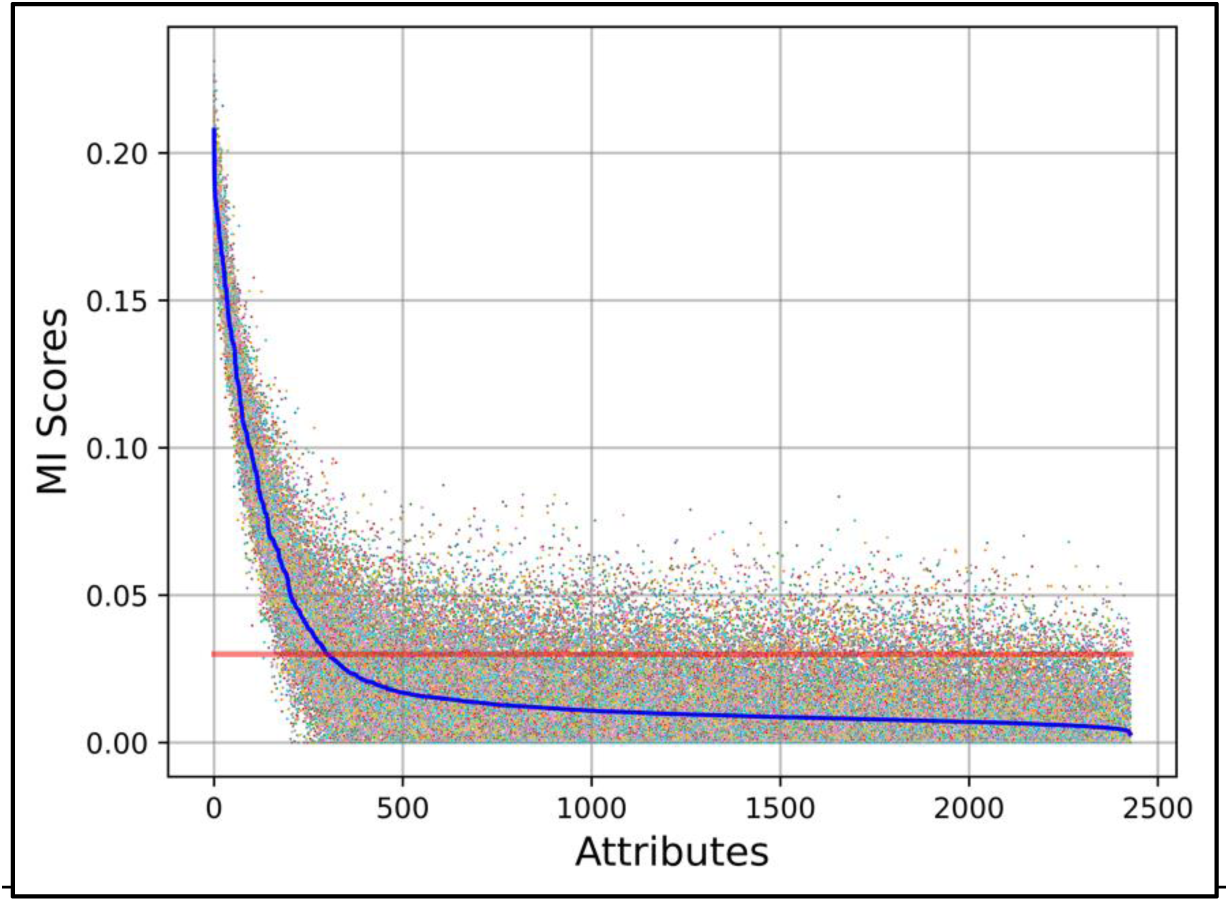
Mutual Information Distribution. The diagram illustrates the values of MI (y-axis) for each peptide attribute (x-axis) from 50 independent trials, the mean <u>*MI*</u> of a feature (blue line), and the threshold value (red line) for feature selection.

The filtered set of features was then assessed for multicollinearity among the features using iterative variance inflation factor (VIF) calculation, with the feature having the highest value (but greater than 5) removed. VIF was recalculated again until the feature with the highest VIF value was less than or equal to 5, a condition that was met, providing us with a list of filtered features. The feature selection methodology was repeated 10 times for increased robustness, resulting in improved feature coverage. After ten runs, we identified a union set comprising 104 features, reducing the original number of features by a factor of about 24. Across these runs, 62 common features were retrieved and further ranked based on descending order of MI value. The ML model development was applied to the set of 104 features. The two-stage feature selection retains only the features that aid in classifying peptides with bioactivity against HCV by removing irrelevant, pairwise collinear, and multicollinear features.

### 2.2 Importance of Features

The features that were common in all ten runs indicate the most essential features according to our feature selection strategy. Figures 3 and 4 show these attributes and their descriptive statistics. Table 1 describes the top 15 features for AHCP recognition, in descending order of MI score.

**Figure 3.**
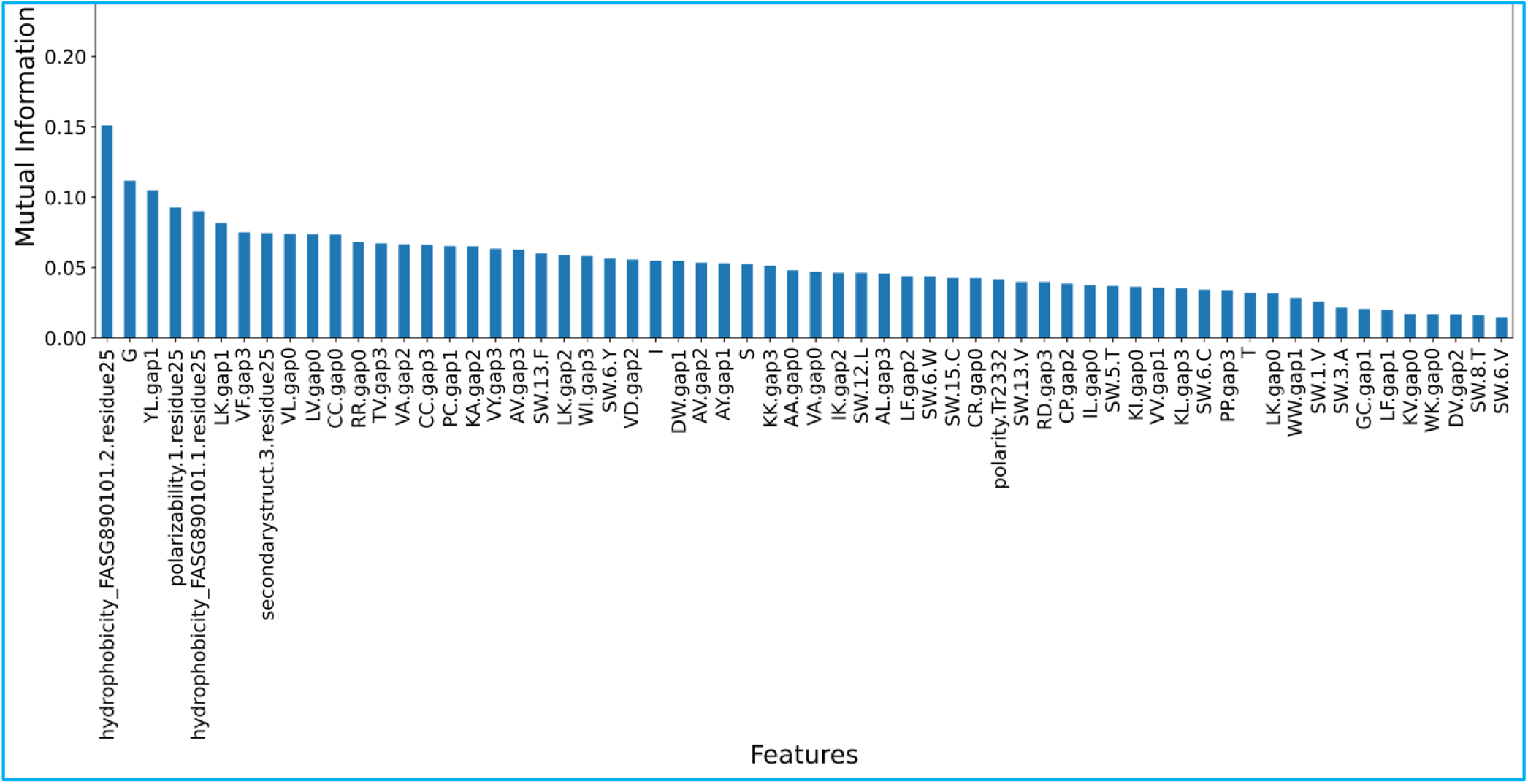
Attribute-specific MI Score. Mean MI scores (y-axis) for peptide features (x-axis) rank-ordered based on MI value.

**Figure 4.**
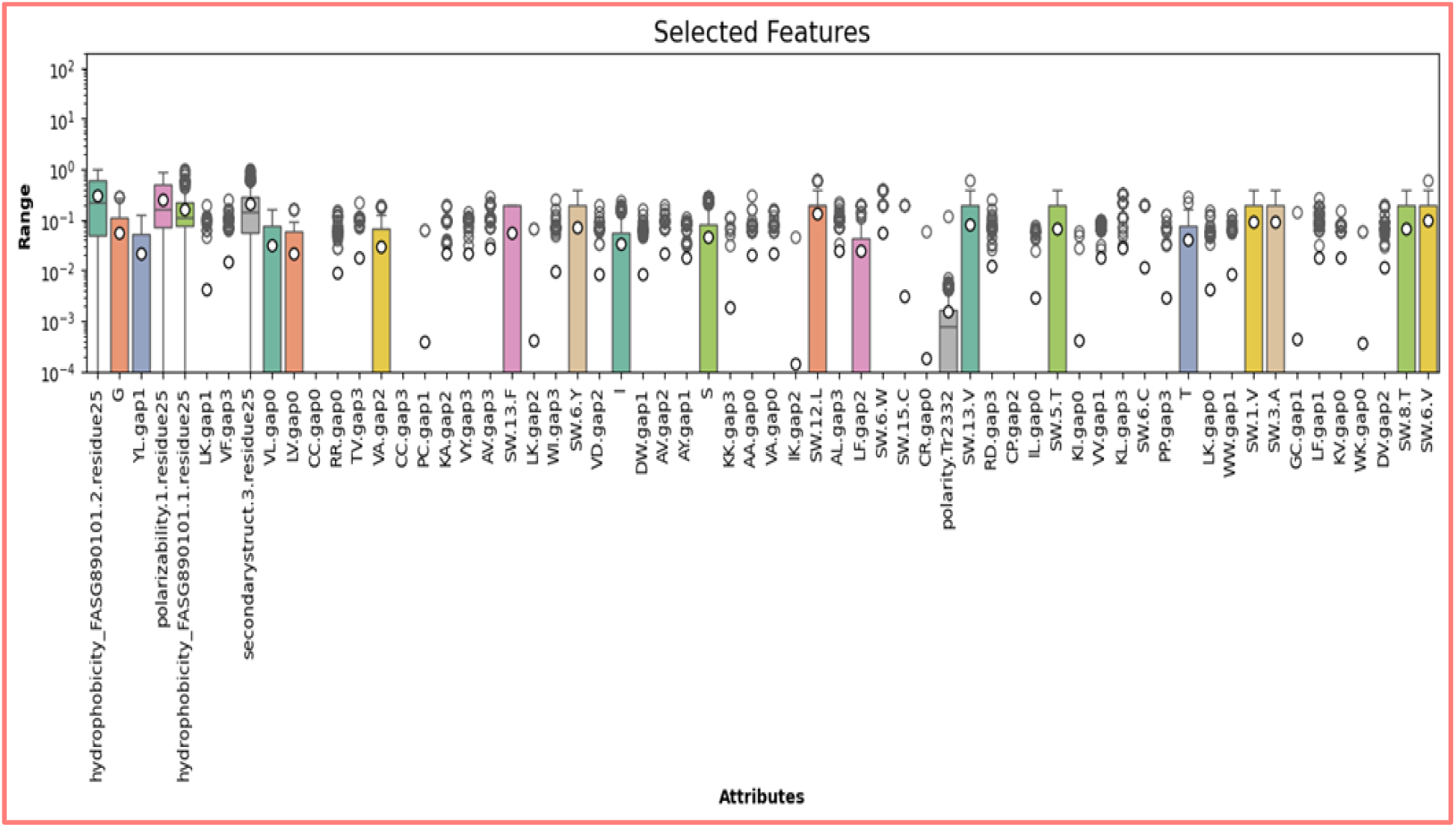
Descriptive Statistics of Peptide Features. Box plots of important features depict their distribution. The data is summarized, showing the minimum, maximum, first, and third quartiles, with the mean represented by an open circle inside each box.

**Table 1.** Selected top 15 features and their description.

| Attributes Acronym | MI score | Feature Description |
| --- | --- | --- |
| hydrophobicity_FASG890101.2.residue25 | 0.151 | Fractional sequence position within which 25% of the residues from group 2 (NTPG) exist. |
| G | 0.112 | Frequency of amino acid G |
| YL.gap1 | 0.105 | Frequency of dipeptide pair Y and L, separated by one residue in between. |
| polarizability.1.residue25 | 0.093 | Fractional sequence position within which 25% of group 1 residues (GASDT that have polarizability 0.000-0.108) exist. |
| hydrophobicity_FASG890101.1.residue25 | 0.090 | Fractional sequence position within which 25% of the residues from group 1 (KERSQD) exist. |
| LK.gap1 | 0.082 | Frequency of dipeptide pair L and K, separated by one residue in between. |
| VF.gap3 | 0.075 | Frequency of dipeptide pair V and F, separated by one residue in between. |
| secondarystruct.3.residue25 | 0.074 | Fractional sequence position within which 25% of the residues forming coil (GNPSD) exist. |
| VL.gap0 | 0.074 | Frequency of dipeptide pair V and L, adjacent to each other. |
| LV.gap0 | 0.074 | Frequency of dipeptide pair L and V, adjacent to each other. |
| CC.gap0 | 0.073 | Frequency of dipeptide pair C and C, adjacent to each other. |
| RR.gap0 | 0.068 | Frequency of dipeptide pair R and R, adjacent to each other. |
| TV.gap3 | 0.067 | Frequency of dipeptide pair T and V, separated by three residues in between. |
| VA.gap2 | 0.067 | Frequency of dipeptide pair V and A, separated by two residues in between. |
| CC.gap3 | 0.066 | Frequency of dipeptide pair C and C, separated by three residues in between. |

According to the feature analysis, the most significant features for AHCPs are the distribution of hydrophobicity corresponding to residues NPTG, frequency of Gly, dipeptide composition of residues YxL, distribution of polarizability corresponding to residues GASDT, and distribution of hydrophobicity corresponding to residues KERSQD. Additionally, dipeptide motifs LXK, VXXXF, VL, LV, CC, RR, TXXXV, VXXXA, CXXXC and distribution of secondary structure elements corresponding to residues GNPSD were selected as other essential features.

How do specific sequence motifs and physicochemical and structural properties assist AHCPs in their function? The heatmap in Figure 5 displays feature values for a sequence, with dendrograms indicating the clustering of sequences with similar attribute profiles. These profiles are significant since it is possible to discern target-specific components of the virus. The top features identified from our analysis can be important for AHCPs in various ways. For instance, to function as entry inhibitory peptides, AHCPs must interact with both hydrophobic and polar amino acids of the HCV envelope glycoprotein 1 (E1 glycoprotein) [16]. The distribution of hydrophobic and polar amino acids in AHCPs is also crucial because of motifs containing glycine and cysteine residues at the N′-terminal and C′-terminal transmembrane domains, respectively, of the E1 glycoprotein. These residues are significant for virus assembly and entry[17]. In general, glycine residues likely provide optimal flexibility to AHC peptides that contain bulky hydrophobic side chains, leading to efficient binding and activity[18].

**Figure 5.**
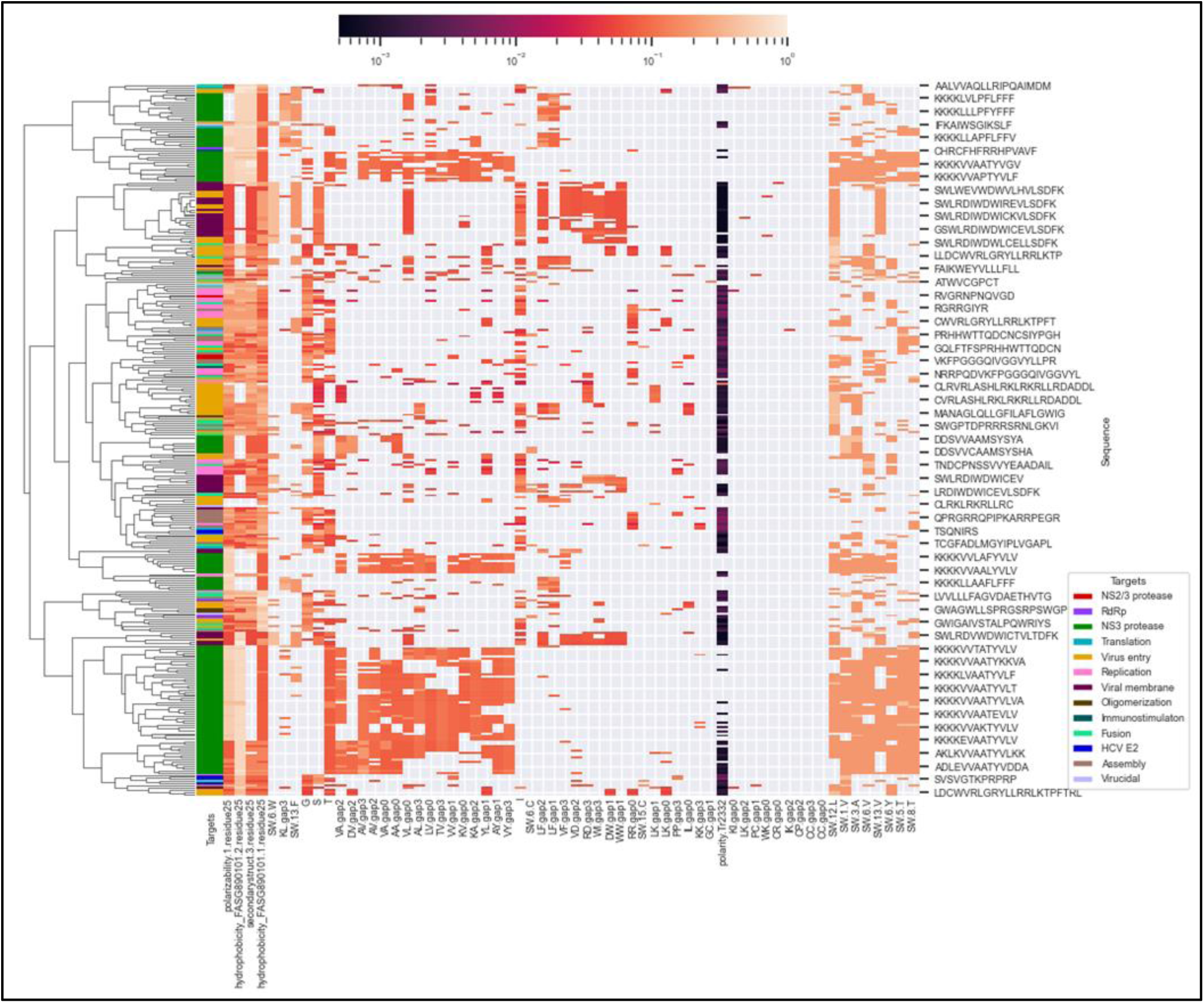
Hierarchical Clustering of anti-hepatitis C peptides dataset. The plot shows peptide features on the horizontal axis (bottom), amino acid sequences on the vertical axis (right), and the peptides’ viral targets on the left.

The distribution of hydrophobic amino acids at the N′-terminal is the leading feature of entry inhibitory peptides that target the pre- or post-binding events of the E1/E2 glycoproteins in HCV. The hydrophobic interactions are crucial for these peptides to work effectively. The CD81 surface receptor binding loops, comprised of the LGAPYWGVF residues, primarily consist of hydrophobic residues[19]. Further, hydrophobicity and positive charge distribution are important features from the point of view of NS3 protease inhibitors, which contain residues DRSHI [20]. Inhibitors targeting *protein translation* are primarily involved in targeting the internal ribosomal entry site (IRES) transacting factors, and amphiphilic peptides with mostly hydrophobic amino acids are shown to bind effectively [21]. HCV assembly relies on the conserved YXXΦ motif, where Φ is a bulky hydrophobic residue, thus explaining the relevance of hydrophobic residues in their AHCP potential [22].

Additionally, the distribution of hydrophobic and positively charged residues are critically important features from the point of view of NS3/4A protease inhibitors, which contain negatively charged aspartic acid residues in addition to serine (polar) and isoleucine (hydrophobic) residues in the active site; thus peptides with optimal distribution of hydrophobic and positively charged amino acid residues would be required for potent AHCPs[23]. Similarly, AHCPs targeting the HCV NS5B, an RNA-dependent RNA polymerase, are needed to bind to two hydrophobic binding pockets predominantly through hydrophobic and polar interactions[24]. The AHCPs targeting the translation and replication process usually bind with human lupus autoantigen (La protein), which binds to the IRES domains and 3’UTR in HCV required for translation and replication or directly binds IRES. In both scenarios, amphiphilic peptides with hydrophobic and charged amino acids were shown to bind effectively[21,25]. The cluster heatmap in Figure 6 provides a visual interpretation of the top-selected features. The selected features can be used to efficiently cluster AHCPs for specific HCV targets and provide insights for further re-engineering and optimisation of peptide candidates. Therefore, critical hydrophobic and polar residue distribution seems vital from the multi-faceted inhibition of HCV entry, replication and assembly, translation, and binding to NS3/4A protease and NS5B polymerase. Thus, the developed ML model comprising of the selected features provides an explanatory framework for an effective prediction and design of AHCPs. Overall, the analysis of the top features selected for AHCPs, and their corresponding targets reveals the molecular basis of their selection from thousands of features initially considered for building the ML models. This underscores the importance of the interpretability of data-driven modelling, uncovering the complex structure–property relationships of AHCPs, which can be useful for designing and re-engineering these peptides.

**Figure 6.**
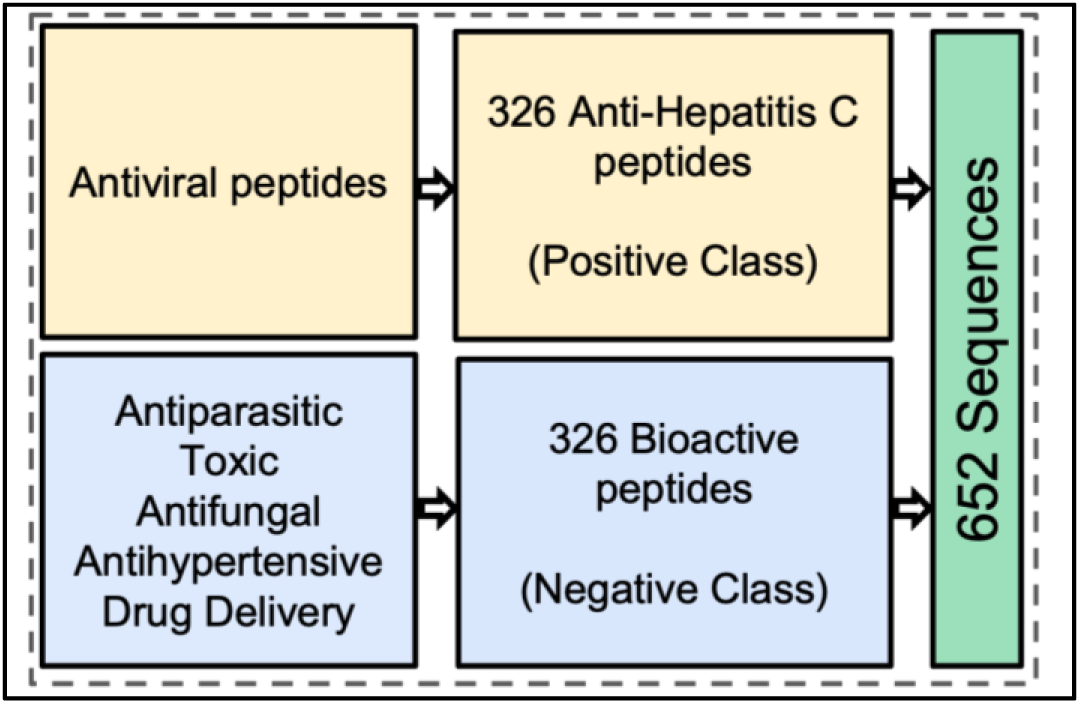
Overview of positive and negative datasets of peptide sequences used in the study.

### 2.3 ML Model Evaluation

We have employed multiple relevant machine learning-based classification algorithms and assessed their potential in identifying AHCPs; see Table 2. We chose these classical ML algorithms based on literature surveys [26] highlighting important ML algorithms from a bioinformatics perspective. After tuning the hyperparameters, we trained and evaluated these models. Table 2 shows the accuracy, precision, recall, specificity, AUROC, and MCC of the six machine learning models on a test dataset consisting of 20% of peptide sequences and an (independent) validation dataset of 18 peptide sequences.

**Table 2.** Performance Evaluation of Machine Learning Algorithms.

| Classifier | Test Accuracy | Test Precision | Test Recall | Test Specificity | Test AUROC | Test MCC | Test F1 Score | Validation Accuracy |
| --- | --- | --- | --- | --- | --- | --- | --- | --- |
| Random Forest | 0.90 | 0.85 | 0.95 | 0.86 | 0.98 | 0.81 | 0.90 | 0.89 |
| Logistic Regression | 0.82 | 0.75 | 0.93 | 0.73 | 0.93 | 0.67 | 0.83 | 0.78 |
| Support Vector Machine (Polynomial) | 0.88 | 0.83 | 0.93 | 0.83 | 0.92 | 0.76 | 0.88 | 0.56 |
| Support Vector Machine (rbf) | 0.84 | 0.78 | 0.92 | 0.77 | 0.94 | 0.69 | 0.84 | 0.5 |
| Support Vector Machine (Linear) | 0.84 | 0.78 | 0.92 | 0.77 | 0.93 | 0.69 | 0.84 | 0.61 |
| Support Vector Machine (Sigmoid) | 0.68 | 0.65 | 0.69 | 0.67 | 0.72 | 0.36 | 0.67 | 0.5 |
| Gaussian Naive Bayes | 0.85 | 0.78 | 0.97 | 0.76 | 0.89 | 0.73 | 0.86 | 0.61 |

The Random Forest classifier demonstrates the highest test accuracy (0.90) and AUROC (0.98) among the models listed. It also maintains high precision (0.85) and recall (0.95) scores for predicting AHCPs. Other notable performers include Gaussian Naive Bayes, with high recall (0.97) and accuracy (0.85), and Support Vector Machine with polynomial kernel, with precision (0.83) and recall (0.93). However, it has a lower MCC (0.76) and validation accuracy (0.56) than Random Forest. Logistic Regression and Support Vector Machines with linear kernels are seen to generally fall behind Random Forest in accuracy and AUROC. We compare the accuracy of the developed model with some of the previous general-purpose antiviral peptide prediction models (Table S1). Despite using a much smaller number of sequences and features, the performance metrics of our models were either similar or better.

## 3. CONCLUSION

We have developed ML models to predict HCV-specific peptides based on their sequence features. In contrast to conventional feature selection, we adopted a twofold method to extract relevant features while eliminating multicollinear variables. The first step involved employing MI to identify relevant features, and in the second step, VIF was used to remove multicollinear variables. After testing various ML algorithms on the reduced dataset, the random forest model provided the best predictive ability and accuracy. The leading features selected for AHCPs and their corresponding targets offer insights into the molecular interactions underpinning their selection from the vast array of features initially evaluated for constructing the ML models. The elucidation of the selected top features provides greater confidence in the developed ML model for AHCP classification. Moreover, the top features can serve as inputs for the targeted design of novel AHCPs and modulation of existing AHCPs. This task is not feasible with previously developed non-specific ML models for antiviral peptides. To our knowledge, this study presents a strategy customised specifically for predicting AHCPs, with the potential to extend its application to designing other classes of antiviral peptides against specific viruses. The ML model is now available on a user-friendly web server, simplifying the process of predicting AHCPs.

## 4. MATERIALS AND METHODS

### 4.1 Preparation of Dataset

We have curated a dataset of peptide sequences that specifically target elements of the Hepatitis C virus; refer to Figure 6. This dataset serves as the primary resource for developing machine learning-based classification models tailored to AHCP bioactivity. The peptide sequences in this dataset have been screened from the AVPdb[11] repository of experimentally validated bioactive antiviral peptide sequences. From this repository, natural peptides with bioactivity against HepC have been selected and enriched further using a filtration scheme. The filtering involves the selection of peptide sequences of length less than 50 residues, followed by removing sequences containing modified, ambiguous or chemically modified amino acid residues. The length criterion is based on the standard definition of *small* linear peptides. Modified residues with chemical moieties such as acetyl or methyl group and ambiguous residues such as norleucine or B = D or N, J = I or L, X = unknown, Z = E or Q, have been removed to eliminate the difficulty in extracting the relevant features classes from such modified residues. Potential duplicate sequences were also removed during the processing step. This resulted in 326 peptide sequences denoting the positive bioactive peptide class against Hepatitis C.

Creating robust and unbiased negative datasets of peptide sequences is essential for improving the predictive capability of ML models. For this study, we selected the sequences from a collection of bioactive peptides with no known antiviral bioactivity[27] to construct an overall balanced negative dataset. These peptides exhibit a range of bioactivities, including antiparasitic, toxic, antifungal, drug delivery, and antihypertensive properties. Peptide sequences longer than 50 residues and those with ambiguous and/or modified residues were dropped, following a similar filtering process as the positive dataset. This resulted in a total of 326 peptide sequences to represent the class of non-AHCPs (but also non-antiviral bioactive peptides). When creating the dataset, we prioritized uniformity in sequence lengths between positive and negative peptide datasets while minimizing redundancy. This involved manually aligning the length distributions of peptide sequences in both sets. The distributions of sequence length and amino acids in the dataset obtained from the post-processing are described in Figure 7.

**Figure 7.**
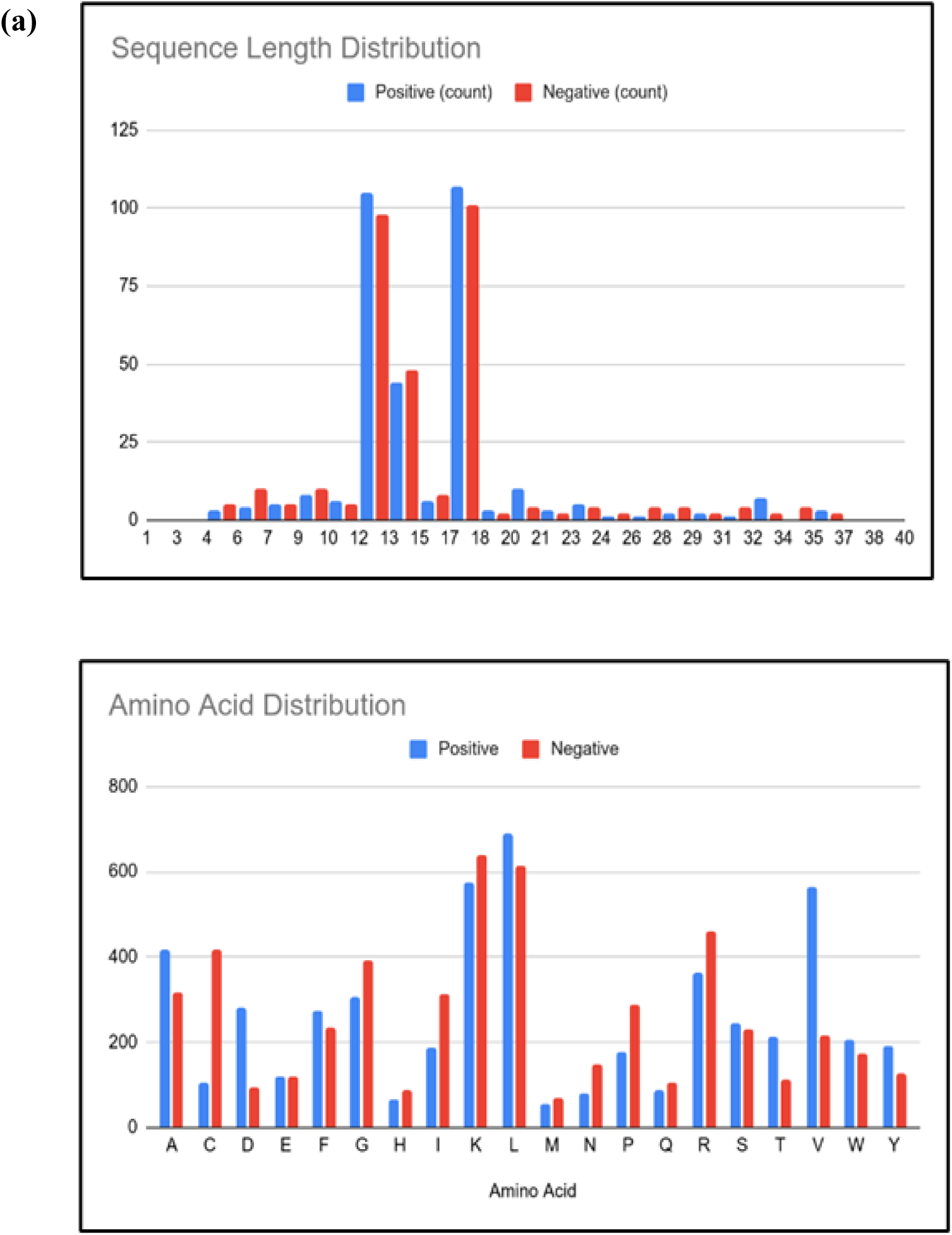
Histograms of Peptide Sequence Dataset for AHCP Classification. (a) Sequence length, and (b) Amino acids.

As we focus on a specific virus, we have created an independent *validation* dataset comprising 18 representative sequences, equally divided between positive and negative classes. The peptides in the positive class have been identified from literature and protein databases. They are known to exhibit AHCP bioactivity, while the peptides in the negative class consist of bioactive peptides without documented antiviral bioactivity.

### 4.2 Feature Descriptors

It is challenging to develop a sequence-based predictive model using only sequence information to rationalize the biological activities of peptides. Hence, sequence features that include composition and physicochemical properties of amino acids are required for robust predictive and inferential abilities of ML classification models. These diverse feature groups create a large multi-dimensional space for ML algorithm activity prediction. For AHCP classification, we computed four distinct groups of features. These features aim to capture global and position-specific (or local) attributes, such as composition and physicochemical properties of the amino acids, and how they are distributed throughout a peptide sequence from its N′-terminal to C′-terminal. The data vectors and computation of the feature groups are defined and discussed in the following subsections.

#### 4.2.1 Amino Acid Composition (AAC)

This feature group contains the relative frequencies of amino acids that comprise a peptide sequence. It is defined as a 20-dimensional array, each element representing one of the 20 amino acids; refer to Figure 8(a). AAC has been commonly employed for classification models such as nuclear receptors[28] and anticancer peptides[29].

**Figure 8:**
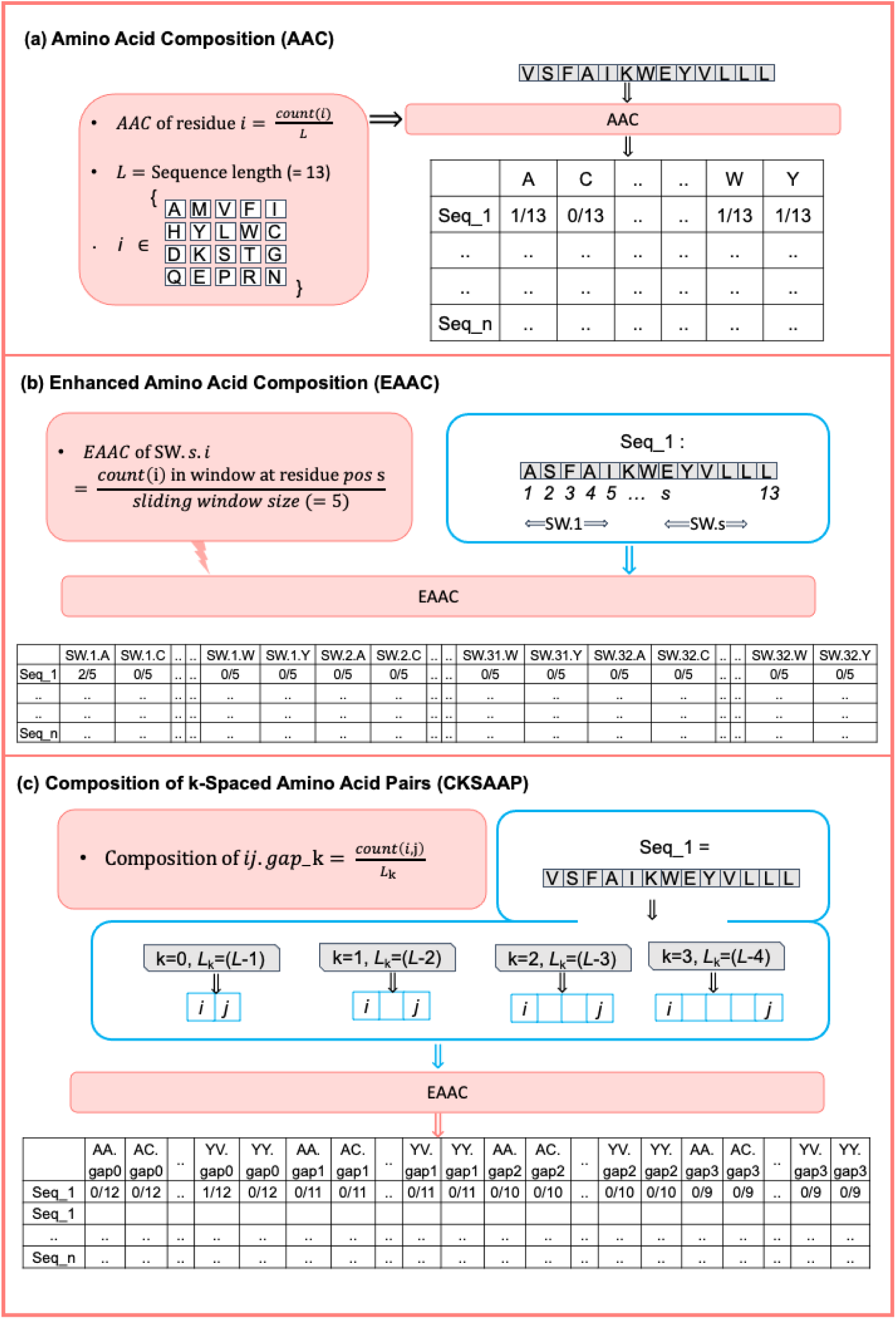
**(a) Amino Acid Composition (AAC).** The calculation of AAC for Seq_1 is described with *i* representing the set of amino acids and N representing the sequence length. For n sequences, the final n×20 array representation of AAC is shown. **(b) Enhanced Amino Acid Composition (EAAC).** It describes the calculation of EAAC for Seq_1. SW.1, SW.2, …, and SW.32 represent the different sequence windows used for AAC calculation. With a maximum length of 36 residues, the total sliding windows are 32. The final n × (32 * 20) array representation of EAAC is shown. **(c) Composition of k-spaced Amino Acid Pairs (CKSAAP).** It describes the calculation of CKSAAP of Seq_1, with *i* and *j* representing the set of amino acids. Note that 400 combinations are possible for each k. The final n × (4 * 400) array representation of CKSAAP for k = 0, 1, 2, 3 is shown

#### 4.2.2 Enhanced Amino Acid Composition (EAAC)

This feature represents position-specific amino acid composition within a sequence window (of a fixed length of 5 contiguous residues) that slides from the N′ to C′ terminal and calculates AAC in each frame, giving a numeric vector; refer to Figure 8(b). The EAAC feature has been successfully applied to predict protein malonylation [30] and lysine crotonylation sites[31].

#### 4.2.3 Composition of k-spaced Amino Acid Pairs (CKSAAP)

This feature represents the probability of a pair of amino acids separated by k spaces or residues where k = 0, 1, 2, 3; refer to Figure 8(c). It attempts to identify the presence of short linear motifs in peptides important for bioactivity. CKSAAP has found utility in predicting membrane protein classes [24], protein crystallization[25], flexible/rigid regions in proteins[26], and structural classes of proteins[32].

#### 4.2.4 CTD encoding

This feature encoding combines three characteristics: composition (C), transition (T), and distribution (D)—each related to the physicochemical properties of amino acids [12,33]. Here, the sequence information of peptides is represented in terms of amino acid properties such as hydrophobicity, polarity, polarizability, charge, secondary structure content, normalized van der Waals volume and solvent accessibility. The amino acids have been further categorized into three subclasses for each property using different labels or indices; see Table 4. Applying CTD encodings has enabled the prediction of protein folding class[34,35], enzyme families[36], RNA-binding proteins[37], protein structures[38] and anti-cancer peptides[29].

**Table 4.** Physicochemical properties and their subdivisions associated with amino acids [35]

| Attribute | Discretised Values |  |  |
| --- | --- | --- | --- |
|  | G1 | G2 | G3 |
| Charge | Positive:<br>KR | Neutral:<br>ANCQGHILMFPSTWYV | Negative:<br>DE |
| Hydrophobicity_CASG920101 | Polar:<br>KDEQPSRNTG | Neutral:<br>AHYMLV | Hydrophobicity:<br>FIWC |
| Hydrophobicity_FASG890101 | Polar:<br>KERSQD | Neutral:<br>NTPG | Hydrophobicity:<br>AYHWVMFLIC |
| Normalised van der Waals volume | Volume range:<br>0-2.78<br>GASTPD | Volume range:<br>2.95-4.0<br>NVEQIL | Volume range:<br>4.03-8.08<br>MHKFRYW |
| Polarity | Polarity value:<br>4.9-6.2<br>RKEDQN | Polarity value:<br>8.0-9.2<br>GASTPHY | Polarity value:<br>10.4-13.0<br>CLVIMFW |
| Polarizability | Polarizability<br>value:<br>0.0-0.108<br>GASDT | Polarizability value: 0.128-0.186<br>CPNVEQIL | Polarizability<br>value: 0.219-0.409<br>KMHFRYW |
| Secondary structure | Helix:<br>EALMQKRH | Strand:<br>VIYCWFT | Coil:<br>GNPSD |
| Solvent accessibility | Buried:<br>ALFCGIVW | Exposed:<br>PKQEND | Intermediate:<br>MPSTHY |

##### 4.2.4.1 Composition (CTDC)

CTDC denotes the composition of physicochemical properties in a sequence. Figure 9(a) shows that if we consider the amino acid *charge* as a property, CTDC calculates the probability of observing *positive*, *negative*, and *neutral* amino acids in the sequence. Considering 8 property types, we have a 24-dimensional vector representation of CTDC.

**Figure 9:**
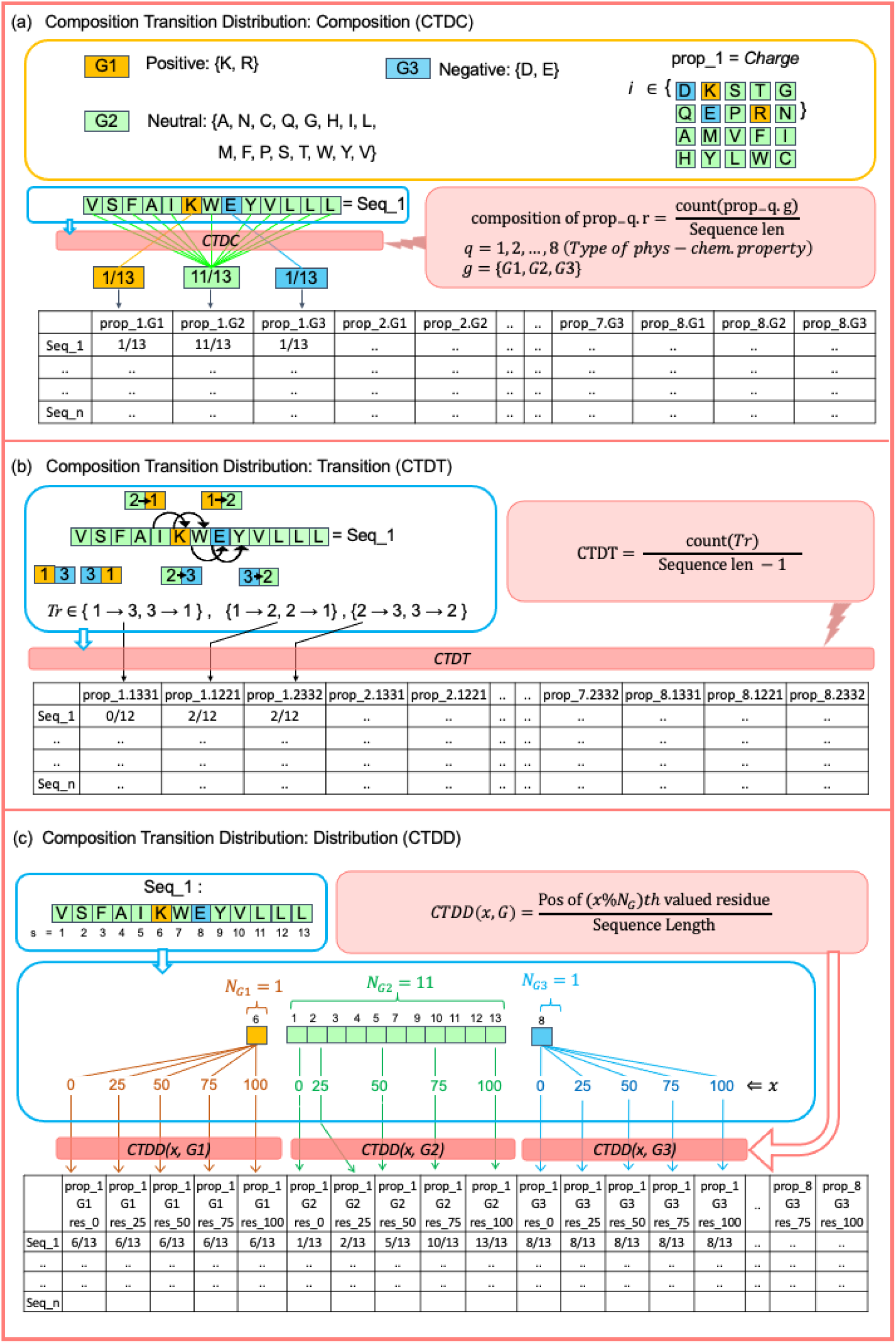
Illustration of Composition, Transition, and Distribution Features. **(a)** Composition (CTDC). It calculates the composition of *positive*, *negative*, and *neutral* amino acid residue classes (represented by colours). Consider prop_1 as charge and N representing the sequence length. The final n × (8 * 3) array is constructed to represent CTDC for the three classes of eight physicochemical properties given in Table 4. (b) Transition–CTDT. In this example, taking charge as property prop_1, the CTDT variable captures the number of transitions of contiguous charged residue classes, encoded by colours, between consecutive residues. The final n × (8 * 3) array is developed to represent transitions for the three classes of eight physicochemical properties. **(c).** Distribution– CTDD. The normalized position relative to the N′ terminal of a peptide sequence VSFAIKWEYVLLL within which the first residue (or 0%), 25%, 50%, 75% and 100% of amino acid residues of a specific property value *g* [e.g., *positive-G1*, *neutral-G2 or negative-G3*] exist. The CTDD of the property *charge* is captured by a 15-dimensional vector.

##### 4.2.4.2 Transition (CTDT)

CTDT refers to the counts of the transitions in the value of a physicochemical property between two consecutive residues when traversing a sequence from the N′ to C′ terminal; see Figure 9(b). For the three discrete values a property can assume, there are three possible transitions (*Tr*), *viz.,* 12 or 21, 13 or 31, and 23 or 32, with 1, 2 and 3 representing the amino acid property values. Considering the 8 property types, we have a 24-dimensional vector representation of CTDT.

##### 4.2.4.3 Distribution (CTDD)

CTDD refers to the distribution of residue physicochemical property (e.g., *charge*) in terms of its values (e.g., whether *positive*, *negative*, or *neutral*) along the sequence length; see Figure 9(c). CTDD of a property is encoded by a five-dimensional vector, where each element is the residue position (normalized by sequence length), starting from N′ terminal, within which the first, 25%, 50%, 75% and 100% of the residues of that specific property-value exist. Considering 8 physicochemical properties, each comprising three nominal values (e.g., *positive*, *negative, and neutral* for a property such as *charge*), we have a fixed 120-dimensional vector representation of CTDD.

### 4.3 Feature Calculation for Peptide Sequences

Python scripts were written for calculating the peptide sequence features— AAC, CKSAAP, EAAC and CTD—given in Figures 8 and 9. The array embeddings shown in these figures were concatenated row-wise to construct the AHCP classification dataset. The features developed were verified using the *ilearnplus* module[39]. The dataset consisted of 2428 attributes, 652 peptide sequence instances of positive and negative classes, and an independent validation set of an additional 18 sequences, which was not used in model training, was compiled; refer to SI.

### 4.4 Feature Selection Methodology

In machine learning, researchers employ different feature selection techniques to enhance models’ generalizability to new data [40]. Mutual information has been used to detect antimicrobial peptides; see Tripathi et al. [41]. In this study, we have computed mutual information [13,14] followed by a variance inflation factor [26] estimation for feature selection. The combination is important because it enhances the dataset by discarding redundancy and collinearity. MI is a widely used concept from information theory to select the most relevant features from a dataset. It is assessed by measuring the dependency of the input features and target variables. Being model-neutral, MI represents a measure of the amount of information that two random variables share. When the MI score is zero or extremely low, e.g., 0.01, it indicates little or no relationship between the feature and the target. Higher values indicate higher dependency.

Multicollinearity arises when two or more independent variables are highly correlated, suggesting redundancy in the information they collectively provide about the dependent variable. The goal is to eliminate multicollinear variables without affecting the model’s quality but rather enrich the dataset. VIF measures the extent of multicollinearity in a set of independent variables and usually quantifies the severity of multicollinearity in an ordinary least squares regression analysis. A high VIF denotes a higher degree of collinearity between the associated independent variables in the model. The combination of MI and VIF for feature selection allowed us to select non-redundant features and eliminate multicollinear features that may lead to models with low confidence and generalizability.

#### 4.4.1 Mutual Information

MI values were calculated using “mutual_info_classif” of the *sklearn* python library [42]. 500 instances of MI calculations were performed, and MI values were then averaged for each feature. Plotting MI values vs attribute indices presented an elbow at <u>*MI*</u>=0.03, and <u>*MI*</u>>0.03 was chosen as a threshold for filtering relevant features. This led us to approximately 200 features from a set of 2428 features.

#### 4.4.2 Variance Inflation Factor

Following the application of the MI filter to select the feature set, an iterative calculation of the variance inflation factor (VIF) [26] was performed on the chosen features. It involves removing the feature with the highest VIF value and then recalculating the VIF of the remaining set of features. This process continues until the highest VIF value is less than or equal to 5. Around 100 features were extracted and later utilized for developing ML classifiers.

The MI and VIF filters effectively extract relevant features. However, due to the randomness in the MI filter, the feature selection method has been repeated tenfold to improve feature coverage. This resulted in identifying 104 relevant features, of which 62 were common across all runs.

### 4.5 Classification Models and Performance Evaluation Metrics

To predict anti-hepatitis C peptides, we have chosen six classical statistical learning algorithms— Random Forest[43,44], Logistic Regression [45,46], SVM [4,47], and Gaussian Naïve Bayes [48,49].

The performance of the models is evaluated using the following metrics and the confusion matrix of predicted classes, providing the number of true positive (TP), true negative (TN), false negative (FN), and false positive (FP) sample predictions.

**Accuracy score (ACC)**

It is used to determine the correctly classified instances out of the total number of instances, given as the ratio of the sum of true positives (TP) and true negatives (TN) versus the sum of all predictions comprising of TP, TN, false negatives (FN) and false positives (FP).

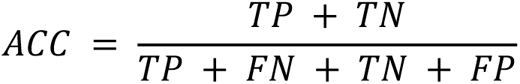

**Precision score (PRE)**

It determines the accuracy of positive class predictions as a ratio of true positives versus the sum of true positives and false positives.

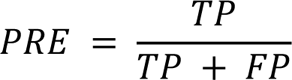

**Recall or Sensitivity (Sn):**

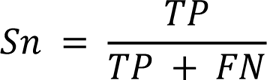

It captures the true positive rate as the ratio of true positives versus the sum of true positives and false negatives.

**Specificity (Sp)**

It measures the model’s ability to identify the negative class correctly. It is the ratio of true negatives versus the sum of true negatives and false positives.

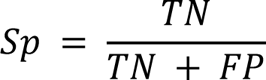

**Area Under ROC Curve (AUROC)**

It tells the probability that a randomly selected AHCP will have a higher predicted probability of being an AHCP than a randomly selected non-AHCP. AUROC > 0.85 is considered excellent performance.

**Matthews Correlation Coefficient (MCC)**

It is a correlation coefficient that achieves a high score only if the classifier correctly predicts the majority of both the positive and negative instances.

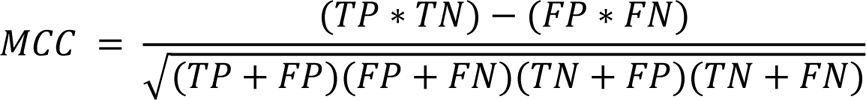

**F1 Score (F1)**

It is expressed as a harmonic mean of precision and recall.

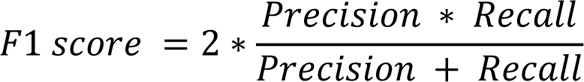

### 4.6 Web Server Development

We have hosted the random forest model on an open-access web server that can be accessed without login requirements. The server’s backend is supported by Flask (version 2.3.3), a web application framework, Pandas (version 2.0.3), and Scikit-learn (version 1.2.2). This web server is hosted on a Linux server that utilises Apache-2 for its operations. The home and prediction result pages of the web server are shown in Figure 13.

**Figure 10:**
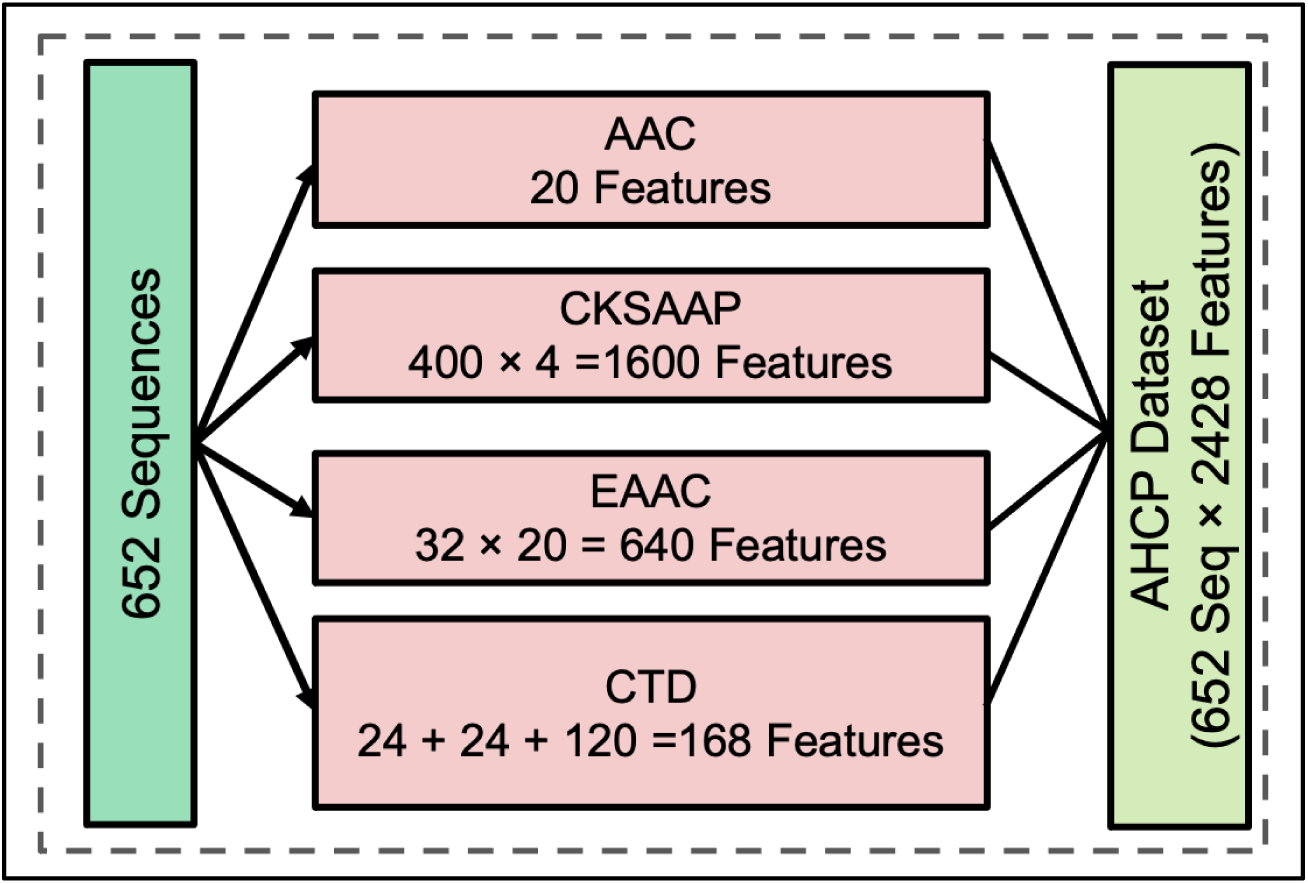
AHCP Dataset. Overview of peptide sequence features.

**Figure 11.**
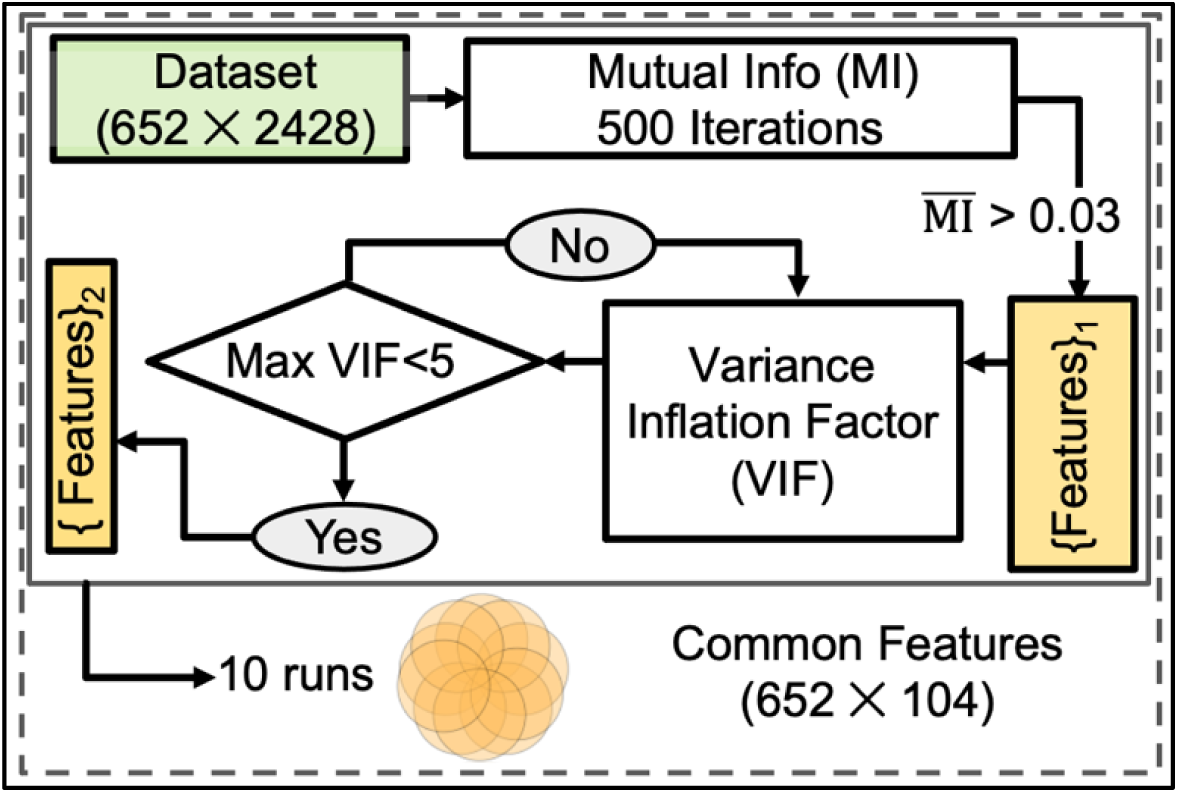
Feature Selection: Schematic representation combining mutual information (MI) and variance inflation factor (VIF).

**Figure 12.**
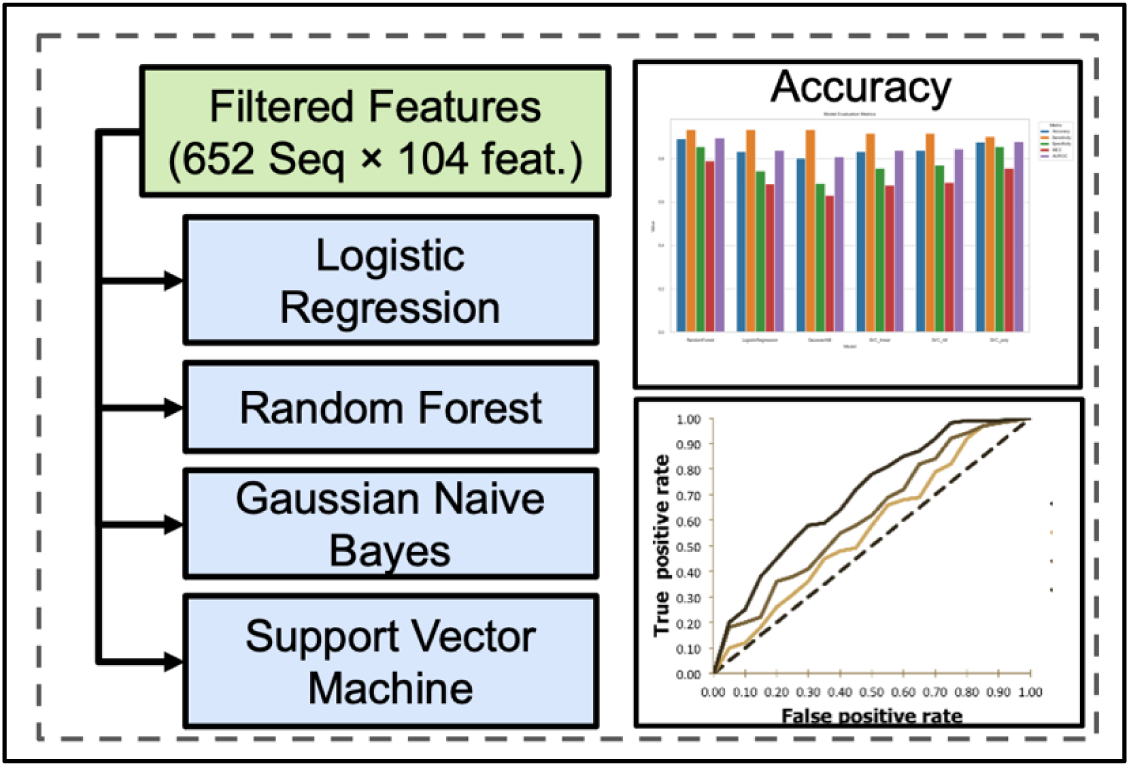
Classification Models: Machine learning algorithms evaluated in the present study.

**Figure 13.**
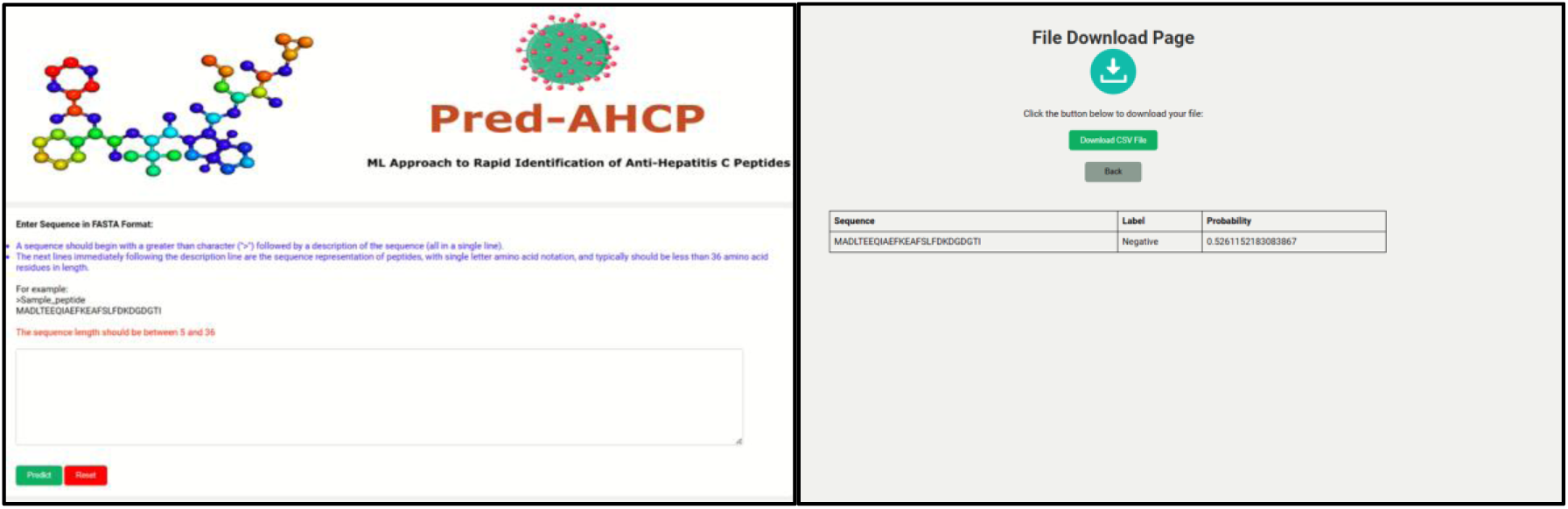
Pred-AHCP Web Server. Snapshots of the Home and Result pages. The input peptide sequences are in FASTA format.

## Abbreviations

HepC: Hepatitis
C HCV: Hepatitis C virus
AHC: Anti-Hepatitis C
AHCPs: Anti-Hepatitis C Peptides
ML: Machine Learning
MI: Mutual Information
VIF: Variance Inflation Factor

## Author Contributions

Akash Saraswat: Data curation; formal analysis; investigation; methodology; validation; writing – original draft.

Utsav Sharma: methodology; software. Aryan Gandotra: web server; software. Lakshit Wasan: web server;

software. Sainithin

Artham: software.

Arijit Maitra: Data curation; formal analysis; investigation; methodology; validation; writing – original draft.

Bipin Singh: Data curation; formal analysis; investigation; methodology; validation; writing – original draft.

## ACKNOWLEDGEMENTS

Akash Saraswat acknowledges the PhD fellowship support from BML Munjal University.

## DATA AVAILABILITY STATEMENT

Link to supporting data: https://github.com/bipiniiith/Pred-AHCP

Web Server URL: http://tinyurl.com/web-Pred-AHCP

## REFERENCES

1. Hepatitis C. [cited 21 Mar 2024]. Available: https://www.who.int/news-room/fact-sheets/detail/hepatitis-c#:~:text=Overview,of%20infection%20without%20any%20treatment.

2. Peptides in chemical space. Medicine in Drug Discovery. 2021;9: 100081.

3. Al-Azzam S, Ding Y, Liu J, Pandya P, Ting JP, Afshar S. Peptides to combat viral infectious diseases. Peptides. 2020;134: 170402.

4. Chowdhury AS, Reehl SM, Kehn-Hall K, Bishop B, Webb-Robertson B-JM. Better understanding and prediction of antiviral peptides through primary and secondary structure feature importance. Sci Rep. 2020;10: 19260.

5. Bin Y, Zhang W, Tang W, Dai R, Li M, Zhu Q, et al. Prediction of Neuropeptides from Sequence Information Using Ensemble Classifier and Hybrid Features. J Proteome Res. 2020;19: 3732–3740.

6. Basith S, Manavalan B, Hwan Shin T, Lee G. Machine intelligence in peptide therapeutics: A next-generation tool for rapid disease screening. Med Res Rev. 2020;40: 1276–1314.

7. Wang G, Vaisman II, van Hoek ML. Machine Learning Prediction of Antimicrobial Peptides. Methods Mol Biol. 2022;2405: 1–37.

8. Tarasova OA, Rudik AV, Ivanov SM, Lagunin AA, Poroikov VV, Filimonov DA. Machine Learning Methods in Antiviral Drug Discovery. Biophysical and Computational Tools in Drug Discovery. 2021; 245–279.

9. Ali F, Kumar H, Alghamdi W, Kateb FA, Alarfaj FK. Recent Advances in Machine Learning-Based Models for Prediction of Antiviral Peptides. Arch Comput Methods Eng. 2023; 1–12.

10. Charoenkwan P, Yana J, Nantasenamat C, Hasan MM, Shoombuatong W. iUmami-SCM: A Novel Sequence-Based Predictor for Prediction and Analysis of Umami Peptides Using a Scoring Card Method with Propensity Scores of Dipeptides. J Chem Inf Model. 2020;60: 6666–6678.

11. Qureshi A, Thakur N, Tandon H, Kumar M. AVPdb: a database of experimentally validated antiviral peptides targeting medically important viruses. Nucleic Acids Res. 2014;42: D1147–53.

12. Govindan G, Nair AS. Composition, Transition and Distribution (CTD) — A dynamic feature for predictions based on hierarchical structure of cellular sorting. 2011 Annual IEEE India Conference. IEEE; 2011. doi:10.1109/indcon.2011.6139332

13. Kraskov A, Stögbauer H, Grassberger P. Estimating mutual information. Phys Rev E Stat Nonlin Soft Matter Phys. 2004;69: 066138.

14. Ross BC. Mutual information between discrete and continuous data sets. PLoS One. 2014;9: e87357.

15. Welcome to. In: Python.org [Internet]. [cited 30 Apr 2024]. Available: https://www.python.org/

16. Hu Z, Rolt A, Hu X, Ma CD, Le DJ, Park SB, et al. Chlorcyclizine Inhibits Viral Fusion of Hepatitis C Virus Entry by Directly Targeting HCV Envelope Glycoprotein 1. Cell Chem Biol. 2020;27: 780–792.e5.

17. Tong Y, Lavillette D, Li Q, Zhong J. Role of Hepatitis C Virus Envelope Glycoprotein E1 in Virus Entry and Assembly. Front Immunol. 2018;9: 1411.

18. Wang J, Chou S, Xu L, Zhu X, Dong N, Shan A, et al. High specific selectivity and Membrane-Active Mechanism of the synthetic centrosymmetric α-helical peptides with Gly-Gly pairs. Sci Rep. 2015;5: 15963.

19. Kumar A, Hossain RA, Yost SA, Bu W, Wang Y, Dearborn AD, et al. Structural insights into hepatitis C virus receptor binding and entry. Nature. 2021;598: 521–525.

20. Zheng F, Yi W, Liu W, Zhu H, Gong P, Pan Z. A positively charged surface patch on the pestivirus NS3 protease module plays an important role in modulating NS3 helicase activity and virus production. Arch Virol. 2021;166: 1633–1642.

21. Dibrov SM, Parsons J, Carnevali M, Zhou S, Rynearson KD, Ding K, et al. Hepatitis C virus translation inhibitors targeting the internal ribosomal entry site. J Med Chem. 2014;57: 1694–1707.

22. Neveu G, Barouch-Bentov R, Ziv-Av A, Gerber D, Jacob Y, Einav S. Identification and targeting of an interaction between a tyrosine motif within hepatitis C virus core protein and AP2M1 essential for viral assembly. PLoS Pathog. 2012;8: e1002845.

23. Meewan I, Zhang X, Roy S, Ballatore C, O’Donoghue AJ, Schooley RT, et al. Discovery of New Inhibitors of Hepatitis C Virus NS3/4A Protease and Its D168A Mutant. ACS Omega. 2019 [cited 13 Sep 2023]. doi:10.1021/acsomega.9b02491

24. Gordon and CP, Keller PA*. Control of Hepatitis C: A Medicinal Chemistry Perspective. 2004 [cited 13 Sep 2023]. doi:10.1021/jm0400101

25. A cell-permeable peptide inhibits hepatitis C virus replication by sequestering IRES transacting factors. Virology. 2009;394: 82–90.

26. James G, Witten D, Hastie T, Tibshirani R. An Introduction to Statistical Learning: with Applications in R. Springer Science & Business Media; 2013.

27. Singh S, Chaudhary K, Dhanda SK, Bhalla S, Usmani SS, Gautam A, et al. SATPdb: a database of structurally annotated therapeutic peptides. Nucleic Acids Res. 2016;44: D1119–26.

28. Bhasin M, Raghava GPS. Classification of nuclear receptors based on amino acid composition and dipeptide composition. J Biol Chem. 2004;279: 23262–23266.

29. Wei L, Zhou C, Chen H, Song J, Su R. ACPred-FL: a sequence-based predictor using effective feature representation to improve the prediction of anti-cancer peptides. Bioinformatics. 2018;34: 4007–4016.

30. Chen Z, He N, Huang Y, Qin WT, Liu X, Li L. Integration of A Deep Learning Classifier with A Random Forest Approach for Predicting Malonylation Sites. Genomics Proteomics Bioinformatics. 2018;16: 451–459.

31. Zhao Y, He N, Chen Z, Li L. Identification of protein lysine crotonylation sites by a deep learning framework with convolutional neural networks. IEEE Access. 2020;8: 14244–14252.

32. Chen K, Kurgan LA, Ruan J. Prediction of protein structural class using novel evolutionary collocation-based sequence representation. J Comput Chem. 2008;29: 1596–1604.

33. Govindan G, Nair AS. Classification of proteins in intracellular and secretory pathway using global descriptors of amino acid sequence. 2011 World Congress on Information and Communication Technologies. IEEE; 2011. doi:10.1109/wict.2011.6141236

34. Dubchak I, Muchnik I, Holbrook SR, Kim SH. Prediction of protein folding class using global description of amino acid sequence. Proc Natl Acad Sci U S A. 1995;92: 8700–8704.

35. Dubchak I, Muchnik I, Mayor C, Dralyuk I, Kim SH. Recognition of a protein fold in the context of the Structural Classification of Proteins (SCOP) classification. Proteins. 1999;35: 401–407.

36. Cai CZ, Han LY, Ji ZL, Chen YZ. Enzyme family classification by support vector machines. Proteins. 2004;55: 66–76.

37. Han LY, Cai CZ, Lo SL, Chung MCM, Chen YZ. Prediction of RNA-binding proteins from primary sequence by a support vector machine approach. RNA. 2004;10: 355–368.

38. Tomii K, Kanehisa M. Analysis of amino acid indices and mutation matrices for sequence comparison and structure prediction of proteins. Protein Eng. 1996;9: 27–36.

39. Chen Z, Zhao P, Li C, Li F, Xiang D, Chen Y-Z, et al. iLearnPlus: a comprehensive and automated machine-learning platform for nucleic acid and protein sequence analysis, prediction and visualization. Nucleic Acids Res. 2021;49: e60–e60.

40. Dhal P, Azad C. A comprehensive survey on feature selection in the various fields of machine learning. Applied Intelligence. 2021;52: 4543–4581.

41. Tripathi V, Tripathi P. Detecting antimicrobial peptides by exploring the mutual information of their sequences. J Biomol Struct Dyn. 2019 [cited 27 Jul 2023]. doi:10.1080/07391102.2019.1695667

42. sklearn.feature_selection.mutual_info_classif. In: scikit-learn [Internet]. [cited 30 Apr 2024]. Available: https://scikit-learn.org/stable/modules/generated/sklearn.feature_selection.mutual_info_classif.html

43. Kumar V, Patiyal S, Dhall A, Sharma N, Raghava GPS. B3Pred: A Random-Forest-Based Method for Predicting and Designing Blood-Brain Barrier Penetrating Peptides. Pharmaceutics. 2021;13. doi:10.3390/pharmaceutics13081237

44. Chang KY, Yang J-R. Analysis and prediction of highly effective antiviral peptides based on random forests. PLoS One. 2013;8: e70166.

45. Park H, Lo-Ciganic W-H, Huang J, Wu Y, Henry L, Peter J, et al. Evaluation of machine learning algorithms for predicting direct-acting antiviral treatment failure among patients with chronic hepatitis C infection. Sci Rep. 2022;12: 18094.

46. Akbar S, Raza A, Zou Q. Deepstacked-AVPs: predicting antiviral peptides using tri-segment evolutionary profile and word embedding based multi-perspective features with deep stacking model. BMC Bioinformatics. 2024;25: 102.

47. Mekni N, Coronnello C, Langer T, Rosa MD, Perricone U. Support Vector Machine as a Supervised Learning for the Prioritization of Novel Potential SARS-CoV-2 Main Protease Inhibitors. Int J Mol Sci. 2021;22. doi:10.3390/ijms22147714

48. Muhammad LJ, Algehyne EA, Usman SS, Ahmad A, Chakraborty C, Mohammed IA. Supervised Machine Learning Models for Prediction of COVID-19 Infection using Epidemiology Dataset. SN Comput Sci. 2021;2: 11.

49. Rui F, Yeo YH, Xu L, Zheng Q, Xu X, Ni W, et al. Development of a machine learning-based model to predict hepatic inflammation in chronic hepatitis B patients with concurrent hepatic steatosis: a cohort study. EClinicalMedicine. 2024;68: 102419.

50. Zhang YP, Zou Q. PPTPP: a novel therapeutic peptide prediction method using physicochemical property encoding and adaptive feature representation learning. Bioinformatics. 2020;36: 3982– 3987.

51. Wei L, Zhou C, Su R, Zou Q. PEPred-Suite: improved and robust prediction of therapeutic peptides using adaptive feature representation learning. Bioinformatics. 2019;35: 4272–4280.

52. Otović E, Njirjak M, Kalafatovic D, Mauša G. Sequential Properties Representation Scheme for Recurrent Neural Network-Based Prediction of Therapeutic Peptides. J Chem Inf Model. 2022;62: 2961–2972.

53. Mahlapuu M, Björn C, Ekblom J. Antimicrobial peptides as therapeutic agents: opportunities and challenges. Crit Rev Biotechnol. 2020;40: 978–992.

54. Charoenkwan P, Anuwongcharoen N, Nantasenamat C, Hasan MM, Shoombuatong W. In Silico Approaches for the Prediction and Analysis of Antiviral Peptides: A Review. Curr Pharm Des. 2021;27: 2180–2188.

55. Schaduangrat N, Nantasenamat C, Prachayasittikul V, Shoombuatong W. Meta-iAVP: A Sequence-Based Meta-Predictor for Improving the Prediction of Antiviral Peptides Using Effective Feature Representation. Int J Mol Sci. 2019;20. doi:10.3390/ijms20225743

56. Qureshi A, Kaur G, Kumar M. AVCpred: an integrated web server for prediction and design of antiviral compounds. Chem Biol Drug Des. 2017;89: 74–83.

57. Nasiri F, Atanaki FF, Behrouzi S, Kavousi K, Bagheri M. CpACpP: Cell-Penetrating Anticancer Peptide Prediction Using a Novel Bioinformatics Framework. ACS Omega. 2021;6: 19846–19859.

58. Hasan MM, Alam MA, Shoombuatong W, Deng H-W, Manavalan B, Kurata H. NeuroPred-FRL: an interpretable prediction model for identifying neuropeptide using feature representation learning. Brief Bioinform. 2021;22. doi:10.1093/bib/bbab167

59. Wang L, Huang C, Wang M, Xue Z, Wang Y. NeuroPred-PLM: an interpretable and robust model for neuropeptide prediction by protein language model. Brief Bioinform. 2023;24. doi:10.1093/bib/bbad077

60. Malik A, Subramaniyam S, Kim C-B, Manavalan B. SortPred: The first machine learning based predictor to identify bacterial sortases and their classes using sequence-derived information. Comput Struct Biotechnol J. 2022;20: 165–174.

61. Khan A, Uddin J, Ali F, Kumar H, Alghamdi W, Ahmad A. AFP-SPTS: An Accurate Prediction of Antifreeze Proteins Using Sequential and Pseudo-Tri-Slicing Evolutionary Features with an Extremely Randomized Tree. J Chem Inf Model. 2023;63: 826–834.

62. Dai R, Zhang W, Tang W, Wynendaele E, Zhu Q, Bin Y, et al. BBPpred: Sequence-Based Prediction of Blood-Brain Barrier Peptides with Feature Representation Learning and Logistic Regression. J Chem Inf Model. 2021;61: 525–534.

63. Müller AT, Hiss JA, Schneider G. Recurrent Neural Network Model for Constructive Peptide Design. J Chem Inf Model. 2018;58: 472–479.

64. Wei S, Liu C, Du L, Wu B, Zhong J, Tong Y, et al. Identification of a novel class of cyclic penta-peptides against hepatitis C virus as p7 channel blockers. Comput Struct Biotechnol J. 2022;20: 5902–5910.

65. Chen S, Liao Y, Zhao J, Bin Y, Zheng C. PACVP: Prediction of Anti-Coronavirus Peptides Using A Stacking Learning Strategy with Effective Feature Representation. IEEE/ACM Trans Comput Biol Bioinform. 2023;PP. doi:10.1109/TCBB.2023.3238370

66. Vishnepolsky B, Gabrielian A, Rosenthal A, Hurt DE, Tartakovsky M, Managadze G, et al. Predictive Model of Linear Antimicrobial Peptides Active against Gram-Negative Bacteria. J Chem Inf Model. 2018;58: 1141–1151.

67. Teimouri H, Medvedeva A, Kolomeisky AB. Bacteria-Specific Feature Selection for Enhanced Antimicrobial Peptide Activity Predictions Using Machine-Learning Methods. J Chem Inf Model. 2023;63: 1723–1733.

68. Rodrigues CHM, Garg A, Keizer D, Pires DEV, Ascher DB. CSM-peptides: A computational approach to rapid identification of therapeutic peptides. Protein Sci. 2022;31: e4442.

69. Grønning AGB, Kacprowski T, Schéele C. MultiPep: a hierarchical deep learning approach for multi-label classification of peptide bioactivities. Biol Methods Protoc. 2021;6: bpab021.

70. Du Z, Ding X, Xu Y, Li Y. UniDL4BioPep: a universal deep learning architecture for binary classification in peptide bioactivity. Brief Bioinform. 2023;24. doi:10.1093/bib/bbad135

71. Fan H, Yan W, Wang L, Liu J, Bin Y, Xia J. Deep learning-based multi-functional therapeutic peptides prediction with a multi-label focal dice loss function. Bioinformatics. 2023;39. doi:10.1093/bioinformatics/btad334

