## Supplementary Information for "Pred-AHCP: Robust feature selection enabled Sequence-Specific Prediction of Anti-Hepatitis C Peptides via Machine Learning"

I. Rank order feature

As presented in Figure 3, the y-axis has been replaced with a feature’s *rank* based on MI values instead of MI values. The high rank indicates a high $\underline{MI}$ value and vice versa. The horizontal line (red) highlights the selected threshold for feature selection, and the blue line represents the mean rank attained by a feature. In this figure, the relevant features correspond to the segment (solid line, blue) above the threshold (red) and exhibit a small rank variance. Whereas below the threshold are the irrelevant features that also have high variance in rank. This also includes the bottom smear of features having MI values about zero.

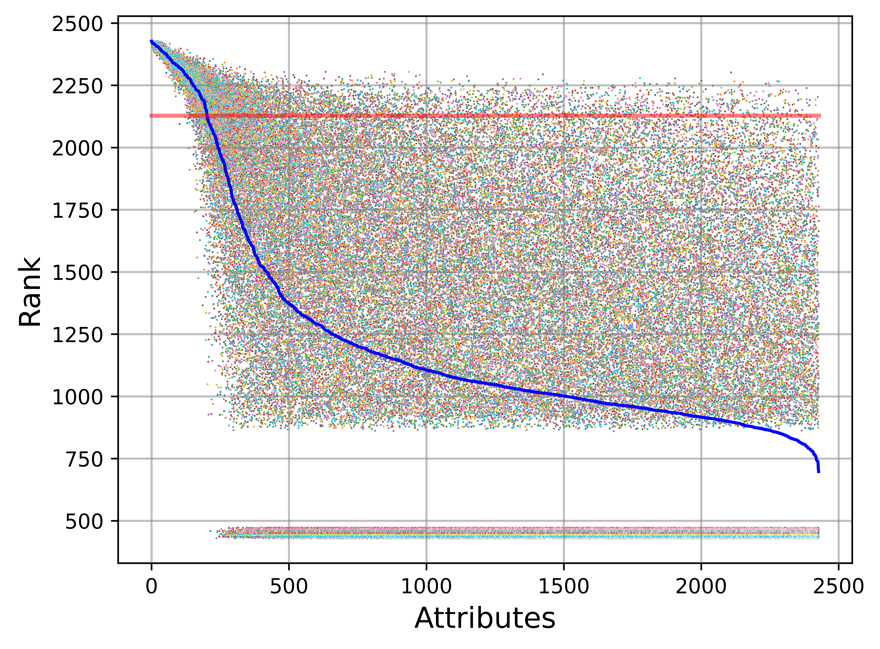

**Figure SS.** **Attribute** **Rank**. The diagram illustrates the rank of attributes based on MI values from 50 independent trials, with the blue curve highlighting the mean rank of a feature and the red line representing the threshold rank.

**II. Application of Machine learning and Deep Learning**

Machine learning and deep learning algorithms have been widely used in the prediction of peptide’s therapeutic bioactivity[[6,50–52]](https://paperpile.com/c/9OdvxC/Jkihg+lk7Lh+YXZ36+E7NZK), antimicrobial[[7,53]](https://paperpile.com/c/9OdvxC/Jr3fG+YC86c) and antiviral[[8,9,54–56]](https://paperpile.com/c/9OdvxC/ibHSC+gZVY5+41ltd+591sn+Zvcaa) bioactivities. Various other bioactivities such as anticancer peptides[[57]](https://paperpile.com/c/9OdvxC/A0a2g), host defence peptides[[52]](https://paperpile.com/c/9OdvxC/E7NZK), neuro-peptides[[5,58,59]](https://paperpile.com/c/9OdvxC/mBTa+NLgTi+FpABz), bacterial sortase enzyme[[60]](https://paperpile.com/c/9OdvxC/jTy5y), antifreeze proteins[[61]](https://paperpile.com/c/9OdvxC/rTAFd), blood-brain barrier peptides[[62]](https://paperpile.com/c/9OdvxC/09rgf) have been predicted using machine learning algorithms. These algorithms have also been used for other related applications, such as designing peptides[[63]](https://paperpile.com/c/9OdvxC/iQdxa). However, very few studies have addressed the issue of peptide prediction specifically for viruses [[64,65]](https://paperpile.com/c/9OdvxC/K0Q2Q+pPVsV) or bacteria [[66,67]](https://paperpile.com/c/9OdvxC/jPiUI+x1fnL). Further, the features necessary for observing specific bioactivities are often not addressed[[10]](https://paperpile.com/c/9OdvxC/fvHsN) despite targeting multiple bioactivities[[68–71]](https://paperpile.com/c/9OdvxC/KJzpJ+KHn1F+e3gm0+cMNcC).

**III. GitHub repositories link**

<https://github.com/bipiniiith/Pred-AHCP>
